# Repeat associated mechanisms of genome evolution and function revealed by the *Mus caroli* and *Mus pahari* genomes

**DOI:** 10.1101/158659

**Authors:** David Thybert, Maša Roller, Fábio C.P. Navarro, Ian Fiddes, Ian Streeter, Christine Feig, David Martin-Galvez, Mikhail Kolmogorov, Václav Janoušek, Wasiu Akanni, Bronwen Aken, Sarah Aldridge, Varshith Chakrapani, William Chow, Laura Clarke, Carla Cummins, Anthony Doran, Matthew Dunn, Leo Goodstadt, Kerstin Howe, Matthew Howell, Ambre-Aurore Josselin, Robert C. Karn, Christina M. Laukaitis, Lilue Jingtao, Fergal Martin, Matthieu Muffato, Michael A. Quail, Cristina Sisu, Mario Stanke, Klara Stefflova, Cock Van Oosterhout, Frederic Veyrunes, Ben Ward, Fengtang Yang, Golbahar Yazdanifar, Amonida Zadissa, David Adams, Alvis Brazma, Mark Gerstein, Benedict Paten, Son Pham, Thomas Keane, Duncan T Odom, Paul Flicek

**Author notes:** Corresponding authors: DTO; PF.

## Abstract

Understanding the mechanisms driving lineage-specific evolution in both primates and rodents has been hindered by the lack of sister clades with a similar phylogenetic structure having high-quality genome assemblies. Here, we have created chromosome-level assemblies of the *Mus caroli* and *Mus pahari* genomes. Together with the *Mus musculus* and *Rattus norvegicus* genomes, this set of rodent genomes is similar in divergence times to the Hominidae (human-chimpanzee-gorilla-orangutan). By comparing the evolutionary dynamics between the Muridae and Hominidae, we identified punctate events of chromosome reshuffling that shaped the ancestral karyotype of *Mus musculus* and *Mus caroli* between 3 to 6 MYA, but that are absent in the Hominidae. In fact, Hominidae show between four-and seven-fold lower rates of nucleotide change and feature turnover in both neutral and functional sequences suggesting an underlying coherence to the Muridae acceleration. Our system of matched, high-quality genome assemblies revealed how specific classes of repeats can play lineage-specific roles in related species. For example, recent LINE activity has remodeled protein-coding loci to a greater extent across the Muridae than the Hominidae, with functional consequences at the species level such as reproductive isolation. Furthermore, we charted a Muridae-specific retrotransposon expansion at unprecedented resolution, revealing how a single nucleotide mutation transformed a specific SINE element into an active CTCF binding site carrier specifically in *Mus caroli*. This process resulted in thousands of novel, species-specific CTCF binding sites. Our results demonstrate that the comparison of matched phylogenetic sets of genomes will be an increasingly powerful strategy for understanding mammalian biology.

## INTRODUCTION

One of the justifications for sequencing many mammalian genomes is to compare these with each other to gain insight into core mammalian functions and map lineage-specific biology. For example, the discovery of human accelerated regions, including the HAR1 gene linked to brain development, relied on comparison between the human and chimpanzee genomes (Pollard et al. 2006). Across the mammalian clade the choice of species to be sequenced and their relative priority have been based on a combination of factors including their value as model organisms (Mouse Genome Sequencing et al. 2002; Gibbs et al. 2004; Lindblad-Toh et al. 2005) or agriculture species (Bovine Genome et al. 2009; Groenen et al. 2012) as well as the value for comparative genome analysis (Lindblad-Toh et al. 2005; Lindblad-Toh et al. 2011). Despite the extreme popularity of mouse and rat as mammalian models, there have been few efforts to sequence the genomes of other closely related rodent species even though greater understanding their specific biology would almost certainly enhance their value as models.

Comparing genome sequences identifies both novel and conserved loci likely to be responsible for core biological functions (Lindblad-Toh et al. 2005), phenotypic differences (Atanur et al. 2013; Liu et al. 2014; Foote et al. 2015), and many other lineage-specific characteristics (Kim et al. 2011; Wu et al. 2014; Foote et al. 2015). Indeed, evolutionary comparisons have even enabled the identification of genomic variation, such as repeat expansions, which can explain aspects of genome and karyotype evolution (Carbone et al. 2014).

Even closely related species can exhibit large-scale structural changes ranging from lineage-specific retrotransposon insertions to karyotype differences. The mechanisms driving these changes may vary between mammalian lineages and the reasons for these differences remain mostly unknown. For example, the rate of chromosomal rearrangement in mammals can vary dramatically between lineages: murid rodents have a rate that has been estimated to be between three times and hundreds of times faster than in primates (Murphy et al. 2005) (Capilla et al. 2016). Transposable elements and segmental duplications have often been found enriched in the vicinity of chromosomal break points (Bailey et al. 2004; Bovine Genome et al. 2009; Carbone et al. 2014). It is not clear whether these transposable elements directly cause chromosomal rearrangement by triggering non-allelic homologous recombination (NAHR) (Janousek et al. 2013) or if they indirectly act via factors such as chromatin structure or epigenetic features (Capilla et al. 2016).

Transposable elements typically make up 40% of a mammalian genome, have variable activity across lineages, and thus can evolutionarily and functionally shape genome structure (Kirkness et al. 2003; Ray et al. 2007). Retrotransposons have numerous links to novel lineage-specific function (Kunarso et al. 2010; Irie et al. 2016). For instance, pregnancy in placental mammals may have been shaped by an increase of activity of the MER20 retrotransposon, which has rewired the gene regulatory network of the endometrium (Lynch et al. 2011). Furthermore, Alu elements have expanded several times in primates with the largest event occurring around 55MYA (Batzer and Deininger 2002) while SINE B2 elements widely expanded in Muroid rodents (Kass et al. 1997) Retrotransposons can affect gene expression by altering pre-mRNA splicing (Lin et al. 2008) or regulatory networks (Jacques et al. 2013; Chuong et al. 2016). For example, lineage-specific transposons can carry binding sites for regulators including the repressor NRSF/REST (Mortazavi et al. 2006; Johnson et al. 2007) and CTCF (Bourque et al. 2008; Schmidt et al. 2012).

The rate of fixation of single nucleotide mutation can also change between different mammalian lineages, for example rodents have a faster rate than primates (Mouse Genome Sequencing et al. 2002). One likely explanation is the shorter generation time observed in rodents compared to primates (Li and Tanimura 1987), (Li et al. 1996). In this hypothesis, most single nucleotide mutations occur during DNA replication in the male germ line and the larger number of passages associated with the rodent’s shorter generation time accumulates more mutations in the same period of time (Goetting-Minesky and Makova 2006).

Thus far, the dynamics of genome evolution between mammalian lineages have been mainly studied mainly by comparing distant genomes (Gibbs et al. 2004; Murphy et al. 2005; Bovine Genome et al. 2009; Lindblad-Toh et al. 2011; Foote et al. 2015), and less frequently using closely-related species(Carbone et al. 2014; Capilla et al. 2016). Comparing distantly-related species can lead to poor resolution of genome structural changes and an inability to assess mechanisms or initial drivers of change. This is due in part to incomplete or uncertain alignments between distant genomes and the inability to unravel multiple evolutionary events that may have occurred in a single genomic region.

At present, primates are one of, if not the only mammalian clade with enough sequenced genomes (Chimpanzee and Analysis 2005; Rhesus Macaque Genome et al. 2007; Locke et al. 2011; Scally et al. 2012; Carbone et al. 2014; Gordon et al. 2016) to facilitate high-resolution studies of genome evolution within a single mammalian lineage (Marques-Bonet et al. 2009; Gazave et al. 2011; Navarro and Galante 2015), (Schwalie et al. 2013). It remains uncertain whether the evolutionary dynamics observed in the primates are common across other mammalian clades.

In this study, we generated high-quality genome assemblies for both *Mus caroli* and *Mus pahari* to create a sister clade for comparison with primate genome evolution. The combination of the *Mus caroli* and *Mus pahari* genomes with the reference mouse and rat genomes mirror, in divergence time and phylogenetic structure, the four Hominidae species with sequenced genomes (human, chimp, gorilla, orangutan). Here we directly compare the processes of genome sequence evolution active within Hominidae and Muridae as two representative clades of mammals.

## RESULTS

### Sequencing, assembly, and annotation of Mus caroli and Mus pahari genomes

We sequenced the genomes of *Mus caroli* and *Mus pahari* females using a strategy combining overlapping Illumina paired-end and long mate-pair libraries with OpGen optical maps (**Figure S1.1A, Methods SM1.1-4**). First, scaffolds were created with ALLPATHS-LG (Gnerre et al. 2011) from the overlapping and 3 kb Illumina mate-pair libraries and then were coupled to the OpGen optical maps to yield 3,079 (*Mus caroli*) and 2,944 (*Mus pahari*) super-scaffolds with a N50 of 4.3 Mb and 3.6 Mb, respectively. We reconstructed pseudo-chromosomes by guiding the assembly based on (i) chromosome painting information and (ii) multiple, closely-related genomes, effectively reducing the assembly bias caused by using only a single reference genome (Kolmogorov et al. 2016). We obtained 20 and 24 chromosomes with a total assembled genome size of 2.55 Gb and 2.47 Gb, respectively, for *Mus caroli* and *Mus pahari*. These two genomes have assembly statistics comparable to the available primate genomes, including chimpanzee, gorilla, and orangutan (**Figure S1.1B**).

We generated RNA-seq data from brain, liver, heart, and kidney in *Mus caroli* and *Mus pahari* to annotate the genes using an integration of TransMap (Stanke et al. 2008), AUGUSTUS (Stanke et al. 2006) and AUGUSTUS-CGP (Konig et al. 2016) pipelines (**Methods SM1.7)**. This approach identified 20,323 and 20,029 protein-coding genes and 10,069 and 9,336 non-coding genes, comparable to the mouse and rat reference genomes (**Figure S1.2A**).

The assembled *Mus caroli* and *Mus pahari* genomes have a low nucleotide error rate, estimated as one sequencing error every 25 to 30 kb based on mapping the mate-pair libraries back to the final corresponding genome assemblies (**Figure S1.1C, Methods SM1.14**). Comparison of the optical maps with the final genome assemblies suggests that up to 3,035 and 1,691 genomic segments could be misassembled, representing 2.5% and 3.1% of the *Mus caroli* and *Mus pahari* genomes, respectively (**Figure S1.1D**). To estimate the gene completion of the two assemblies, we inspected the alignment coverage of protein-coding genes conserved across all vertebrates (**Method SM1.15**). The alignment coverage was 93.3% and 93.2% for the *Mus caroli* and *Mus pahari* assemblies respectively, values that fall within the range (91.6% to 94.7%) for corresponding primate genomes (**Figure S1.2B**).

Previous phylogenetic analyses of the *Mus* genus have relied on the sequence of cytochrome b, 12S rRNA and the nuclear *Irbp* gene to broadly estimate a 2.9-7.6 MY divergence among the *Mus caroli*, *Mus pahari*, and *Mus musculus* species (Veyrunes et al. 2005; Chevret et al. 2014). We refined this estimate using the whole genome assemblies to create a complete collection of the four-fold degenerate sites found in amino-acids conserved across mammals. In specific and highly conserved amino acids, the third base within the coding triplet is thought to be under virtually no selective constraint, meaning neutral rates of change can be estimated by comparing the accumulation of mutations within these sites. We then estimated the divergence time separating *Mus musculus* with *Mus caroli* and *Mus pahari* by anchoring our analysis on a mouse-rat divergence time of 12.5 MY, an estimate based on fossil records (Jacobs and Flynn 2005)(**Figure S1.3A**, **Figure 1A**, **Method SM1.16).**

Our estimates show that *Mus pahari* diverged from the *Mus musculus* lineage 6.0 MYA with a 95% confidence interval ranging from 5.1 to 7.5 MYA and *Mus caroli* diverged 3.0 MYA with a 95% confidence interval ranging from 2.6 to 3.8 MYA (**Figure 1A**). We observed no introgression or incomplete lineage sorting among these four species that could affect the divergence time estimate (**Figure S1.3B**, **Method SM1.17**). These results were robust to (i) the choice of the gene categories from which we selected the four-fold degenerate sites and (ii) the evolutionary model used to make the divergence estimates (**Figure S1.3A, Method S1.16**).

**Figure 1.**
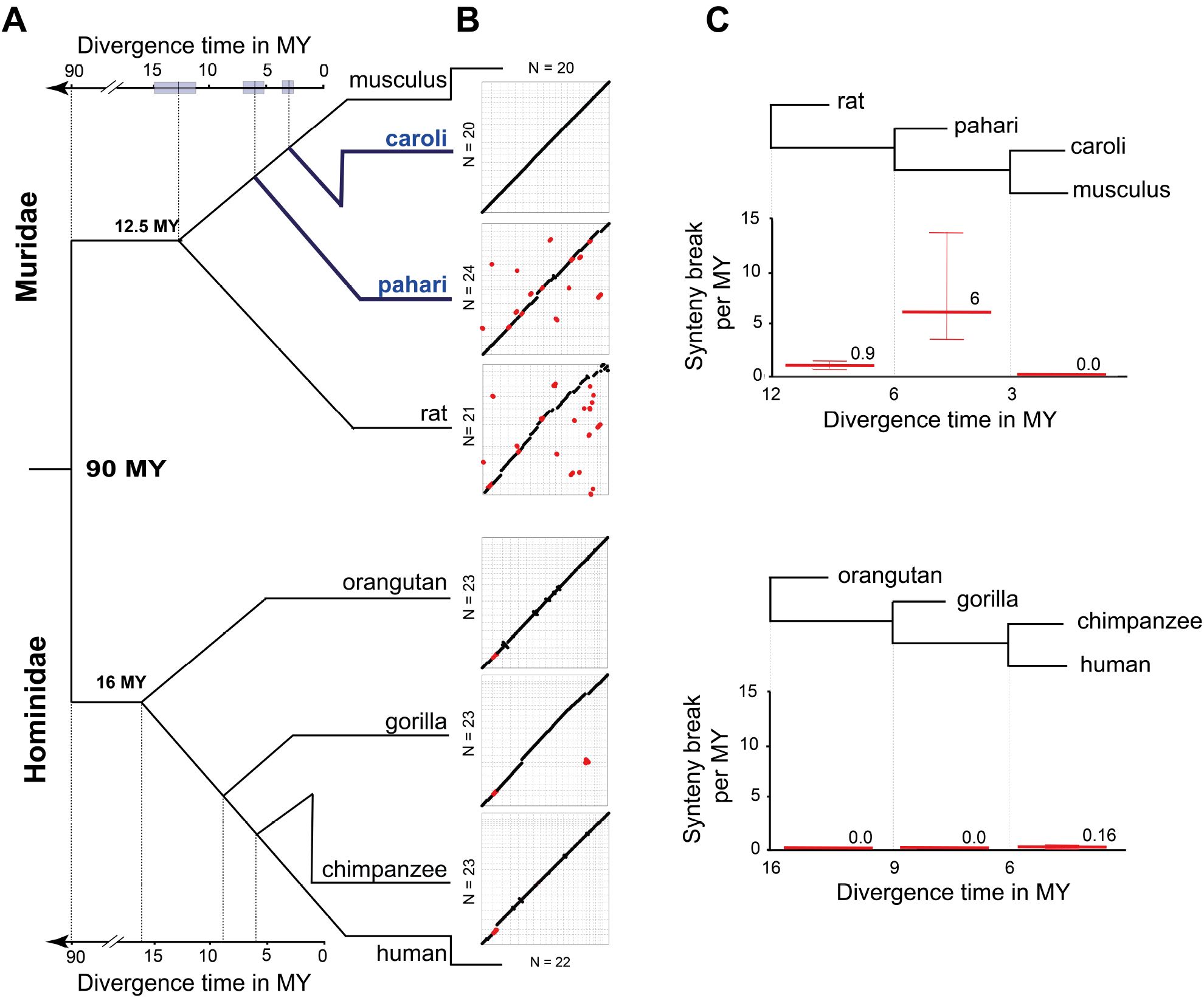
Muridae genomes undergo large chromosomal rearrangements in punctuate bursts, resulting in greater structural diversity than primates. **A** Phylogenetic tree showing that the divergence time of the four Muridae species mirrors that of the four Hominidae species. The *Mus* species in blue were sequenced and assembled for this study. The 95% confidence interval of the divergence time estimation is shown by the shaded boxes (see **Method SM1.16**). **B** Dot plots of whole-genome pairwise comparison between *Mus musculus* and the three other Muridae (top), and between human and the three other Hominidae (bottom). The chromosomes of *Mus musculus* and human were ordered by chromosome number. The chromosomes of the other species were ordered to optimize the contiguity across the diagonal. Red dots represent large (>3 Mb) inter-chromosomal rearrangements (fusion/fission and translocation). **C** The rate of synteny breaks between sequential internal branch points of the Muridae and Hominidae clades. Muridae have undergone a punctuate increase in the rate of syntenic breaks between 3 and 6 MYA.

### A punctuated event of chromosomal rearrangements shaped the Mus musculus and Mus caroli ancestral karyotype

In rodents, chromosome numbers evolve much more rapidly than among most other mammalian clades including primates (Ferguson-Smith and Trifonov 2007). To compare the evolutionary dynamics of large (>3 Mb) inter-chromosomal rearrangements, we performed pairwise whole-genome alignments of the Muridae and Hominidae genomes (**Figure 1B, Figure S1.4**). Hominidae karyotypes, like most mammalian clades, are highly stable, typically showing only one or two unique breaks for each species (Ferguson-Smith and Trifonov 2007) (**Figure S1.4E).**

In contrast, our analysis reveals that the Muridae clade appears to have been subjected to punctate periods of accelerated genome instability interspaced with periods of more typical stability. For example, a period of massive genome rearrangement occurred in the shared ancestor of *Mus caroli* and *Mus musculus* after the split with *Mus pahari* (3-6 MYA) that resulted in 20 synteny breaks found only in *Mus caroli* and *Mus musculus* (**Figure 1C**). Notably, over the most recent 0-3 MY, the karyotypes of *Mus caroli* and *Mus musculus* have been stable with no large genome rearrangements. Second, rat shows 19 lineage-specific synteny breaks when compared with *Mus pahari*, meaning that the rat karyotype more closely resembles that of *Mus pahari* than the karyotypes of the two other *Mus* species. Regardless of whether the rat-specific changes were introduced gradually or in one or more punctuated events, the overall impact on the genome (~20 large breaks) is vastly greater than observed in Hominidae in a roughly corresponding divergence time (orangutan versus human: 1 large break) (**Figure S1.4**).

In order to find a potential molecular mechanism driving the punctate increases of interchromosomal rearrangement, we asked if the inter-chromosomal breakpoints between *Mus musculus* and *Mus pahari* were enriched in repeat elements. Repeat elements are thought to drive chromosome rearrangement by increasing local homology and then inducing NAHR (Hedges and Deininger 2007; Robberecht et al. 2013). We found a significant enrichment of LTR retrotransposons with a concurrent age of the rearrangement events, i.e. 3-6 MY old (Empirical p-value, p < 10^-3^**, Figure S1.5**). We also found an enrichment, although not statistically significant, of SINE elements of the same age. When considering the set of repeats of all ages, there was no observed enrichment at breakpoints for any type of repeat (**Figure S1.5)**. This result suggests that specific LTR repeats may increase local susceptibility to inter-chromosomal rearrangement by NAHR.

In summary, our results detail a punctate event of chromosome reshuffling that happened in the Muridae lineage between 3-6 MYA and that has led to the observed karyotype of laboratory mice. The enrichment of 3-6 MY old LTR elements at the chromosomal breakpoints suggests that this class of retrotransposons is involved in the mechanisms driving these large-scale events.

### Divergence and turnover of genomic sequences and segments are accelerated in Muridae, particularly for LINE retrotransposons

We next tested whether the genome of Muridae evolves faster in than that Hominidae by comparing the rate of nucleotide variation within each clade, and focussing again on the four-fold degenerate sites (**Figure 2A, Method SM 3.1**). We found that the Muridae clade shows six-fold increases in the rate of change within these neutral locations when compared to the Hominidae clade.

**Figure 2.**
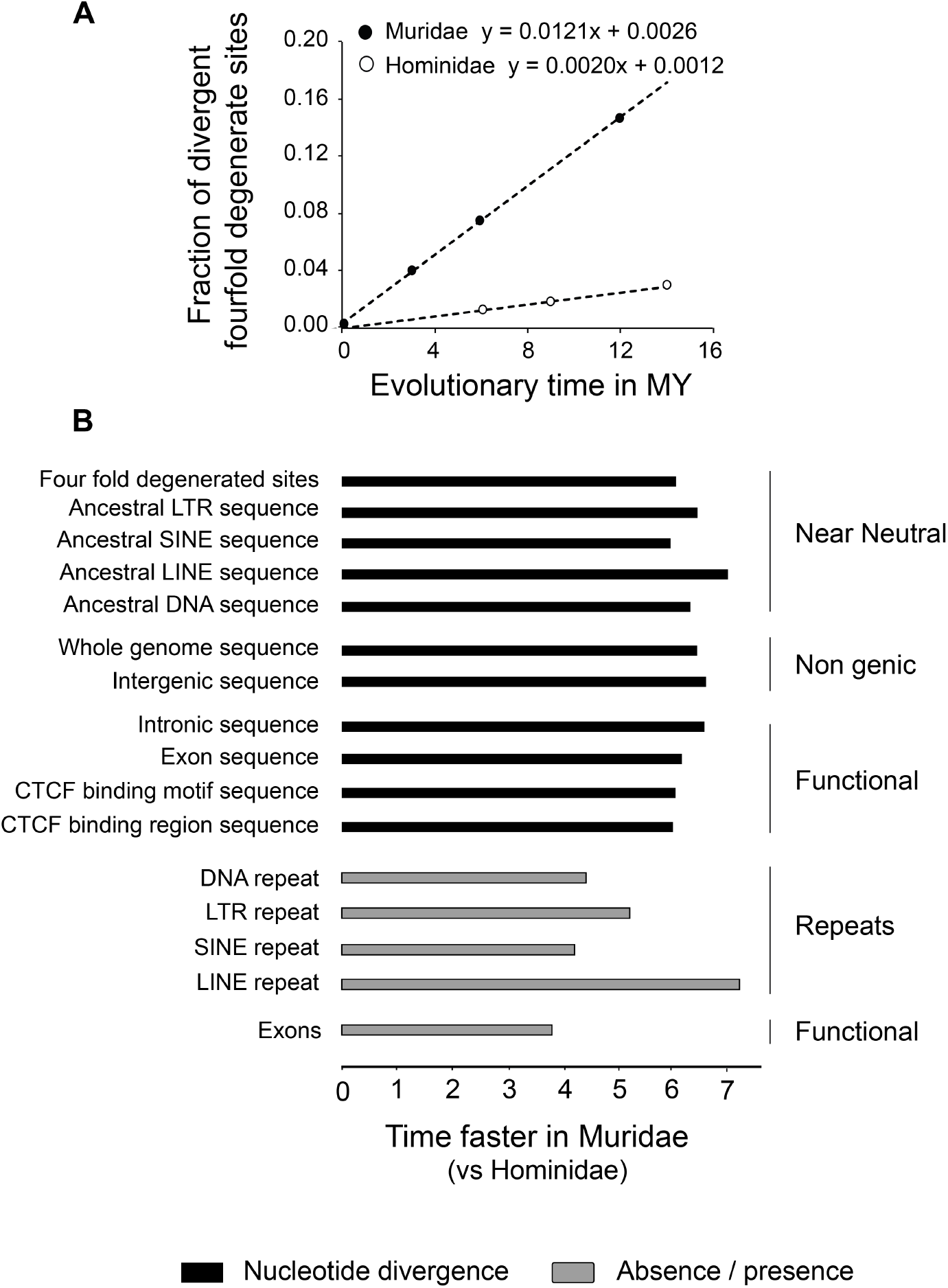
Acceleration of mutational rates in the Muridae lineage. **A** The evolutionary rate calculated from four-fold degenerate sites in Muridae (black) and in Hominidae (white). **B** The bar chart shows the ratios of evolutionary rates between Muridae and Hominidae. Mouse versus human ratios were calculated for rates of nucleotide divergence (black bars) and the turnover rates (grey bars) for specific genomic regions (**Method SM3.2**).

We took a similar approach to establish how rapidly sequence changes occur in the whole genome as well as in specific classes of genomic elements, including ancestral repeats such as LTR, SINE, LINE, and DNA repeats, exons, and CTCF binding motifs (**Figure 2B**). The rate of nucleotide variation change reflects different evolutionary constraints, consistent with Gaffney, et al (Gaffney and Keightley 2006) **(Figure S2.1A)**. Nevertheless, across all inspected categories, Muridae genome evolution is accelerated between six-and seven-fold when compared to primates (**Figure 2B**).

We next quantified how rapidly entire genomic segments are gained and lost among these four rodent species. Similar to nucleotide variation, different types of elements show differing rates of turnover **(Figure S2.1B)**. Because DNA transposons, as opposed to retrotransposons, lost their activity early in the primate and rodent lineages (Mouse Genome Sequencing et al. 2002; Pace and Feschotte 2007) we used the empirically observed turnover of DNA transposons as a background rate. Notably, this background rate of DNA repeat evolution in rodents is approximately four and a half-fold higher than in Hominidae.

In both clades, protein-coding exons are more stable than DNA transposons, as expected. In contrast, both SINE and LTR retrotransposons are actively expanding in a lineage-specific manner and show higher turnover than DNA transposons in both rodents and primates. (**Figure 2B, Figure S2.1B**). Moreover, in both clades, the rates of SINE and LTR element turnover are similar to each other and, when compared to the turnover rate of DNA transposons, exhibit approximately the same relative increase. This suggests that Muridae and Hominidae have a generally comparable activity of SINE and LTR retrotransposons when compared to DNA transposons. However, in Muridae LINE retrotransposons are ~1.5 times more active than LTR and SINE elements, and appear to have greatly accelerated activity when compared to the rate found in primates (ANCOVA, p-val < 10^-3^) (**Figure 2B, Figure S2.1B**).

In summary, our results detail the remarkably rapid evolution of Muridae genomes. Common classes of repeat elements expand between 4.1 and 7.7 fold faster in rodents than in Hominidae genomes. Most notably, LINE retrotransposon activity is highly accelerated in Muridae, and has typically resulted in the birth of several hundred Mb of novel genomic sequence (69-374 Mb) in each assayed rodent genome.

### Accelerated LINE retrotransposon activity has shaped coding gene evolution in rodents

We next asked how retrotransposon activity has changed during the evolutionary history of both clades. We first estimated in each genome the age of every retrotransposon by calculating the sequence identity between the retrotransposon and the consensus sequence, which is an approximation of the ancestral repeat. Since the sequence of transposable elements evolves nearly neutrally, the relationship between the sequence identity and the estimated age of a repeat is approximately linear (**Method SM4.1)** (Liu et al. 2009).

Our analysis confirmed previous reports (Batzer and Deininger 2002) that a major event of SINE Alu element retrotransposition occurred in the primate lineage, peaking at ~55 MYA and subsequently decreasing to the current basal activity (**Figure 3A)**. In contrast, LINE and LTR elements show relatively low but consistent activity during primate evolution (**Figure S3.1B, Figure 3A**). As in primates, LTR elements in rodents also appear to be relatively quiescent over recent evolutionary time. For SINE elements in the Muridae, there has been a consistent level of moderate activity including insertion events from the SINE B2 family previously shown to carry a CTCF binding site (Bourque et al. 2008; Schmidt et al. 2012).

**Figure 3.**
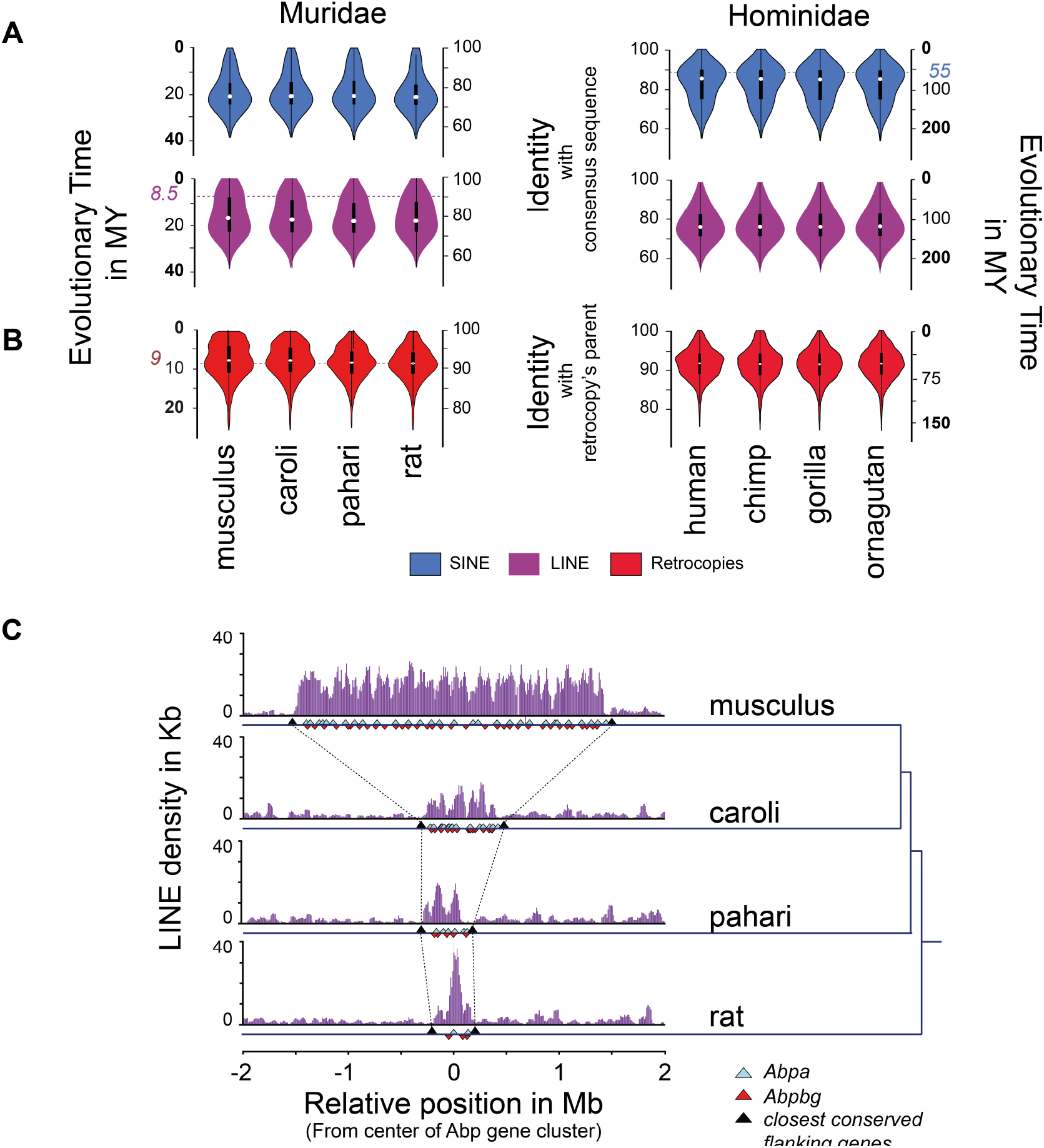
Recent LINE activity can remodel protein-coding gene loci. **A** Identity plots of SINE (blue) and LINE (purple) elements with their repeat consensus for each Muridae (left) and Hominidae (right) species. The age of the transposable elements was estimated using the nucleotide divergence from ancestral SINE and LINE elements (**Methods SM4.1**). The dashed lines indicate the estimated peaks of the most recent expansions in *Mus musculus* and human. **B** Identity plots of retrocopies (red) with their parental genes for each Muridae (left) and Hominidae (right) species. The age of the retrocopies was estimated by the nucleotide divergence from ancestral retrocopies and the corresponding parental genes (**Methods SM4.3**). The dashed line indicates the peak of the most recent expansion in *Mus musculus*. **C** Representation of the density of LINE elements in the *Abp* gene cluster for *Mus musculus*, *Mus caroli, Mus pahari,* and the rat. The blue and red triangles represent the *Abp* genes (*Abpa* (*Scgb1b*) in blue, *Abpbg* (*Scgb2b*) in red), and the black triangles represent the closest flanking genes (upstream: *Scnb1* and downstream: *Uba2*) shared by the four Muridae species.

The most striking difference in retrotransposition activity between the Hominidae and Muridae clades is the greatly accelerated expansion of LINE elements in rodents beginning approximately 8.5 MYA, which has continued at an elevated activity level (**Figure 3A**). This increase has resulted in a substantial enrichment (6-14%; Fisher, p < 10^-16^) of species-specific LINE retrotransposons in all four Muridae species (**Figure S3.1C**).

The LINE-L1 retrotranscriptase machinery can reshape mammalian genomes by capturing RNAs and re-inserting retrotranscribed copies into the genome, as in the case for processed pseudogenes (Esnault et al. 2000). We observed that the rate of mRNA retrocopy integration has increased within the same evolutionary window as the recent LINE expansion in rodents (**Figure 3B**). This increase in mRNA retrotransposition is not found in Hominidae genomes. We also found a small number of chimeric transcripts caused by retrogene insertions in Muridae genomes (**Figure S3.2**, **Methods SM4.4**)

In addition, LINE retrotransposons can act as substrate for NAHR, thus driving copy number variation and gene cluster expansion (Startek et al. 2015; Janousek et al. 2016). The *Secretoglobin* (*Scgb*) gene cluster containing *Scgb1b* and *Scgb2b* genes, also called the *Androgen-binding protein* (*Abp*) gene cluster containing *Abpa* and *Abpbg* genes (Laukaitis et al. 2008) illustrates this effect. *Abp* is involved in mating preference (Laukaitis et al. 2012) and incipient reinforcement in the hybrid zone where the geographic range of two mouse subspecies make secondary contact (Bimova et al. 2011). Since the mouse-rat ancestor, this gene cluster has progressively expanded in the Muridae lineage with the greatest number of copies observed in the *Mus musculus* genome (**Figure 3C**). Importantly, in the four genomes, LINE retrotransposons are enriched within the *Abp* gene cluster compared either with adjacent intergenic regions (Empirical p-value, p < 10^-5^) or with collections of single genes matched for total gene number (Empirical p-value, p < 10^-2^, **Methods SM4.6) (Figure 3C).** LTR elements are also enriched within the *Abp* gene cluster (Empirical p-value, p < 10^-5^, **Figure S3.3).**

Taken together, the dramatic, recent, and still-active expansion of LINE activity in rodents has had important functional consequences for the Muridae genome, ranging from a wave of retrocopy integrations to gene cluster expansions.

### Retrotransposition of SINE B2_Mm1 elements drove a species-specific expansion of CTCF occupancy in Mus caroli

Previous studies have shown that the SINE B2 element carries a CTCF binding motif and can thus drive the expansion of CTCF binding in rodents (Bourque et al. 2008; Schmidt et al. 2012). We took advantage of the closely related Muridae genomes to investigate the molecular mechanisms behind this expansion. We determined the genome-wide binding for CTCF in livers of the four Muridae by performing ChIP-seq experiments (**Figure 4A, Methods SM1.11**). In addition, we used a previously published dataset to identify CTCF genome-wide binding in immortalized lymphoblast cells from four primate species (Schwalie et al. 2013). We found between **~24,000** to **~48,000** CTCF binding sites across the four Muridae species, and between **~21,000** to **~57,000** across the four Hominidae species (**Figure S4.1a**).

**Figure 4.**
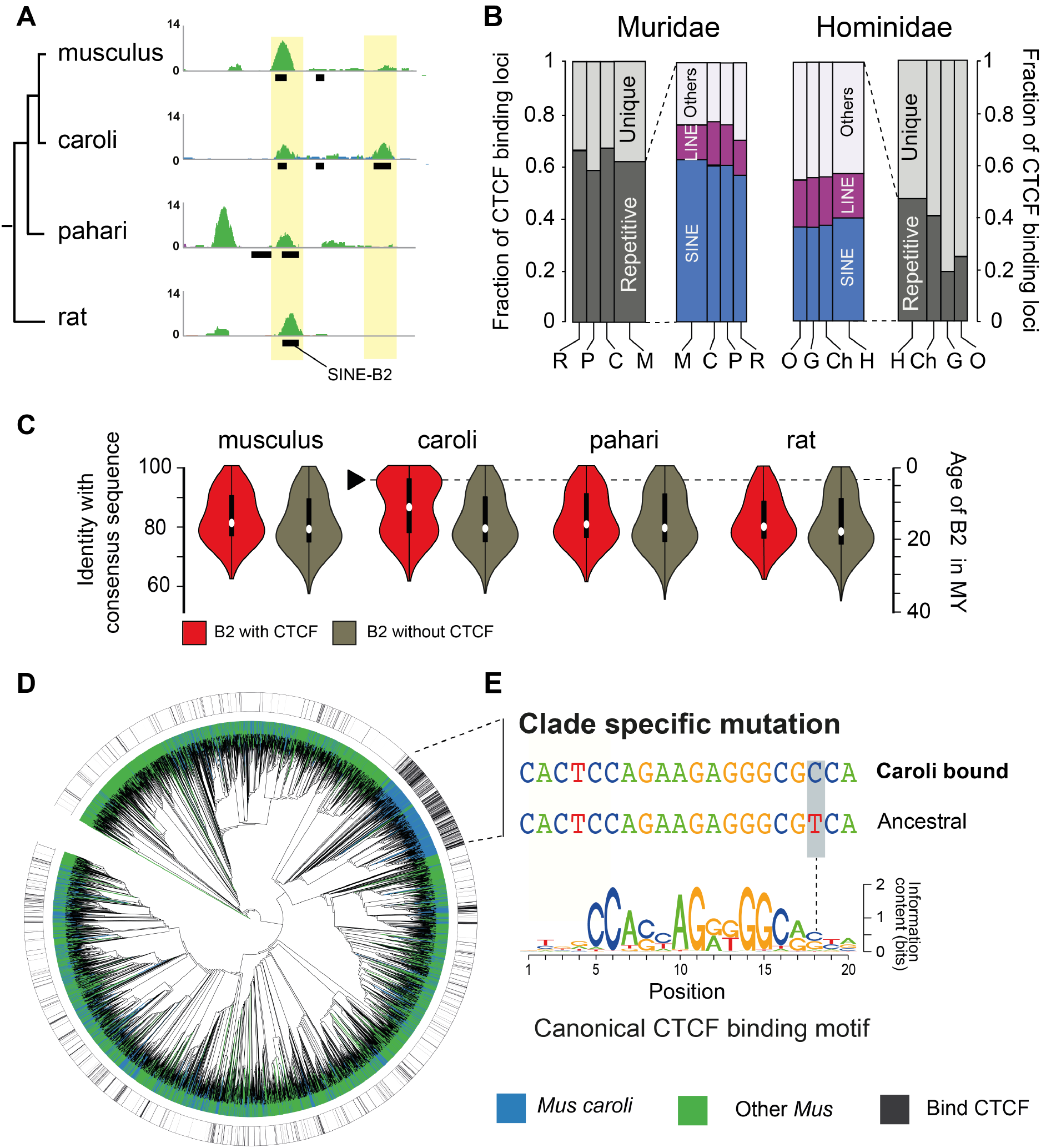
A single nucleotide mutation in a *Mus caroli*-specific expanding SINE B2 element created thousands of novel CTCF binding events. **A** CTCF occupancy in the genome is shown by green tracks. The black squares show the location of SINE B2 retrotransposons. The yellow boxes represents two examples of a SINE B2 occupied by CTCF. **B** Fraction of transposable elements with CTCF binding in both Muridae (left) and Hominidae (right). M=*Mus musculus*; C=*Mus caroli*; P=*Mus pahari*; R=rat; H=human; Ch=chimpanzee; G=gorilla; O=orangutan. **C** Identity plots of SINE B2 with their consensus sequence, either occupied by CTCF (red) or not (brown) (Methods SM4.1). The black arrow indicates a recent wave of SINE B2 expansion carrying CTCF binding sites in *Mus caroli*. **D** Neighbor-joining tree of SINE B2_Mm1 sequences from the three *Mus* species. The blue branches represent sequences from *Mus caroli*. The green branches represent sequences from *Mus musculus* or *Mus pahari*. The black lines in the outside tracks indicates the presence of a CTCF binding event. **E** A single nucleotide variation exists between the ancestral CTCF binding motif carried by the SINE B2_Mm1 element (middle) and a CTCF binding motif (top) carried by the elements recently expanded in *Mus caroli*. This branch specific motif is enriched in CTCF occupancy.

As expected, the CTCF binding sites were overrepresented in SINE retrotransposons in Muridae compared to Hominidae (Fisher, p-val <10^-6^; **Figure 4B**). SINE elements carrying a CTCF binding site were enriched in SINE B2 compared to random expectation (Empirical p-value, p <10^-5^, **Figure S4.1B**). We then asked if any particular mouse species showed enhanced B2 retrotransposition resulting in novel lineage-specific CTCF binding sites. We estimated the age of the B2 elements in the four Muridae species, and found an overrepresentation of young elements positive for CTCF binding in *Mus caroli* (**Figure 4C**). Based on the distribution of repeat ages, this recent wave of CTCF binding site expansion started early in the *Mus caroli* lineage, approximately 3MYA. By comparison, the Hominidae genomes show no similar expansion of CTCF occupancy driven by retrotransposition (**Figure S4.2**).

Next, we asked whether the *Mus caroli*-specific expansion of CTCF binding could be attributed to a particular SINE B2 subfamily. We found an overrepresentation of SINE B2_Mm1 occupied by CTCF specifically in *Mus caroli* when compared with the other rodents (Empirical p-value, p < 10^-5^; **Figure S4.1D**). Among the **20,248** B2_Mm1 elements in *Mus caroli*, 16% (4,151) showed CTCF binding *in vivo*. In contrast, a significantly smaller fraction of B2_Mm1 elements were occupied by CTCF in the other three species of Muridae (2-5%, Fisher test, p < 10^-6^). These results suggest that a B2_Mm1 element carrying an active CTCF binding site has expanded in a species-specific manner in *Mus caroli.*

Notably, the SINE B2_Mm1 family became active specifically in the mouse lineages after the rat-mouse divergence since fewer than 50 B2_Mm1 loci are present in the rat genome. Since the rat-mouse split, B2_Mm1 elements have continued to expand along all three mouse lineages independently, when compared to the ancestral rodent genome. Indeed, we also found a similar overrepresentation of species-specific B2_Mm1 elements in the *Mus musculus* and *Mus pahari* genomes, but these were not associated with a CTCF binding expansion **(Figure S4.3)**.

To understand why CTCF binding loci were expanding only in *Mus caroli*, we determined a B2_Mm1 sequence similarity tree within all three *Mus* species using neighbor joining (**Method SM5.5**). This revealed a monophyletic origin for the majority (59%) of B2_Mm1 elements occupied by CTCF in *Mus caroli* **(Figure 4D)**. This cluster is predominantly composed of *Mus caroli* B2_Mm1 sequences (87%) as well as a handful of B2_Mm1 sequences from the two other *Mus* species. The presence of *Mus musculus* and *Mus pahari* B2_Mm1 sequences suggest that either representatives of this cluster existed, albeit at low copy number, in the ancestral *Mus* species or that there has been random mutation of B2_Mm1 sites in the other lineages.

Sequence analysis suggests that this cluster is enriched in CTCF binding occupancy because of a single nucleotide difference from the ancestral sequence. Specifically, a substitution of a cytosine for a thymine at the position 17 resulted in a more favorable consensus CTCF sequence motif **(Figure 4E)**. Indeed, over 20% of the B2 elements with this motif and not belonging to the B2_Mm1 subfamily were associated with CTCF binding, compared to only 4.7% of the B2 elements with the B2_Mm1 ancestral CTCF motif (Fisher; pval < 10^-16^, **Figure S4.3B**).

In summary, our analysis revealed that a single nucleotide mutation has introduced enhanced CTCF binding affinity into a SINE B2 element present in the *Mus* ancestor. This mutated retrotransposon massively expanded in *Mus caroli* adding more than 2,000 species-specific CTCF binding sites in less than 3MY.

## DISCUSSION

We generated high-quality chromosome-level assemblies of the *Mus caroli* and *Mus pahari* genomes in order to compare the dynamics of genome evolution between the Hominidae and the Muridae. Combining the genomes of *Mus caroli* and *Mus pahari* with those of *Mus musculus* and *Rattus norvegicus* yields a collection of closely related Muridae genomes that are similar in phylogenetic structure and divergence times to Hominidae (human-chimp-gorilla-orangutan). This enables direct comparisons of genome evolutionary dynamics between humans and their most important mammalian models.

Our results provide a detailed description of the remarkably rapid evolution of the Muridae genomes compared to Hominidae within a similar time window. Although the genome-wide increased nucleotide divergence in the Muridae lineage was previously known (Mouse Genome Sequencing et al. 2002), our analysis shows that all categories of genomic annotation and function have similar relative acceleration when compared to Hominidae. The rate change between the two clades is similar, regardless of whether the genomic region is under evolutionary constraint (e.g. coding exons) or apparently evolving neutrally (e.g. ancestral repeats). Thus, the entire genomic system including coding, regulatory and neutral DNA is evolutionary coupled, implying that differences in mutation fixation rate should largely explain the observed acceleration in Muridae.

Although the generation time of Muridae is much shorter than that of Hominidae (Li et al. 1996), this difference alone cannot fully explain the difference between evolutionary rates that we observe. Specifically, wild Muridae have a generation time of approximately 0.5 years (Phifer-Rixey and Nachman 2015), while in Hominidae it is between 20-30 years (Langergraber et al. 2012). This ratio of generation time (40-60) is much higher than the observed ratio of evolutionary rate (6-7), suggesting an important contribution from factors other than generation time (Bromham 2009). We can reduce the effect of generation time by half by considering the increased rate of mutation accumulation per generation in the genome of Hominidae (Uchimura et al. 2015). A further consideration is the effective population size, which is at least one order of magnitude larger in the Muridae compared to the Hominidae (Geraldes et al. 2011; Schrago 2014). Effective population size is a critical parameter to define the mutation fixation rate in a population (Charlesworth 2009). Taken together, we can estimate the effect of population size on the increased mutation fixation rate in Hominidae compared to Muridae to an upper limit of a factor of four. However, considering the complexity of factors influencing the observed evolutionary rate, we cannot exclude other factors such as potential variation in evolutionary rates within the lineage histories that could explain part of these differences.

Our analysis also revealed a different dynamic of karyotype evolution between Muridae and Hominidae. While the Hominidae karyotypes have remained very stable over the past 15 MY (Ferguson-Smith and Trifonov 2007), within a similar period of time Muridae were subject to punctuate periods of accelerated karyotype instability interspaced with periods of more typical stability. These periods of karyotype instability co-occur with specific LTR repeat insertion at chromosomal breakpoints. Our analysis indicates that the rat karyotype is closer to the Murinae ancestor which confirms previous suggestions (Zhao et al. 2004). Several studies suggest that karyotype differentiation is a direct cause of speciation (Kandul et al. 2007; Garagna et al. 2014)). Moreover, a strong link has been made between explosive speciation and periods of karyotype instability in various lineages (Dobigny et al. 2017). In the *Mus* lineage, the *Nannomys* subgenus includes the highest number of species and greatest karyotype diversity (Chevret et al. 2014). Interestingly, the *Nannomys* diverged from the *Mus musculus* lineage between the *Mus caroli* and *Mus pahari* splits (Veyrunes et al. 2005; Veyrunes et al. 2006), i.e. in the same window of increased karyotype instability that we describe here.

Additionally, the analysis of transposable element activity in Muridae and Hominidae has shown that the three main classes of retrotransposons are active in both lineages. This activity has varied over time, and each lineage was subject at some point in their evolutionary history to lineage-specific bursts of retrotransposon activity. For instance, LINE elements had a recent expansive burst specifically in Muridae that is likely still active today. Indeed, the LINE retrotransposon content, even in inbred laboratory mouse strains, can substantially vary (Nellaker et al. 2012). We observed two different functional consequences of repeat-driven lineage-specific genome evolution. First, the progressive expansion of the *Abp* gene cluster across Muridae was correlated with an enrichment of LINE and LTR elements (Janousek et al. 2016). These retrotransposons increase local genome homology and mediate copy number variation via nonallelic homologous recombination (Janousek et al. 2013; Startek et al. 2015), leading to gene expansion. Interestingly the *Abp* gene cluster is involved in mating preference within the peripatric hybrid zone where two mouse subspecies make secondary contact (Bimova et al. 2011). Together, this suggests that transposable elements are involved in the genomic mechanisms driving reproductive isolation between *Mus* sub-species in hybrid zones.

Another observed consequence of repeat driven lineage-specific evolution has been the species-specific expansion of CTCF occupancy sites across the *Mus caroli* genome. Indeed, we demonstrated the effect of a single nucleotide substitution in a SINE B2 followed by expansion of this element to rapidly create thousands of new *Mus caroli*-specific CTCF binding locations. The interplay between nucleotide variation and transposition is a powerful evolutionary mechanism that can disrupt and remodel species-specific regulatory programmes (Kunarso et al. 2010; Schmidt et al. 2012; Mita and Boeke 2016).

We demonstrate that comparing multiple, closely related genomes is one of the most powerful approaches to understand the biology and evolution of a single species. As the number of sequenced genomes rapidly expands in the next ten years (Koepfli et al. 2015), the analysis strategy employed here for the *Mus caroli* and *Mus pahari* genomes and the comparative analysis between Muridae and Homidae can be applied to diverse clades.

## METHODS

### Sequencing and Assembly of Mus caroli and Mus pahari genomes

Genomic DNA was extracted from one *Mus caroli/EiJ* and one *Mus pahari/EiJ* female using Invitrogen’s Easy-DNA kit (K1800-01). 180 bp overlapping paired-end libraries were prepared following Gnerre et al (Gnerre et al. 2011) and 3 kb mate-pair libraries were prepared following

Park et al (Park 2013). These libraries were sequenced using the Illumina HiSeq 2000 platform. The reads were assembled into contigs and scaffolds using the ALLPATHS-LG assembler (Gnerre et al. 2011). High molecular weight DNA was extracted from *Mus caroli/EiJ* and *Mus pahari/EiJ* following the protocol in **Supplementary Material section SM1.2** to construct an optical map using the OpGen platform. The OpGen Genome-Builder software was used to assemble the NGS scaffolds into super scaffolds based on the optical map. Super scaffolds and scaffolds were assembled into pseudo-chromosomes with Ragout (Kolmogorov et al. 2016). To guide the assembly, Ragout used a multiple alignment constructed with Progressive Cactus (Paten et al. 2011). This alignment included the scaffolds of *Mus caroli*, *Mus pahari* and the genomes of *Mus musculus* (C57BL/6NJ GRCm38/mm10 assembly) and *Rattus norvegicus* V5.0. See **Supplementary Material sections SM1.1-SM1.7** for more details.

### Gene annotation

*Mus caroli* and *Mus pahari* genes were annotated using a combination of three annotation pipelines: TransMap (Stanke et al. 2008), AUGUSTUS (Stanke et al. 2006), and a new mode of AUGUSTUS called Comparative AUGUSTUS (AUGUSTUS-CGP) (Konig et al. 2016). The GENCODE set of *Mus musculus* transcripts (M8 release) (Harrow et al. 2012) was used with the TransMap pipeline. In addition, RNA-seq data was used with the AUGUSTUS and AUGUSTUS-CGP pipelines. To prepare the RNA-seq data, RNA was extracted from multiple tissues (brain, liver, heart, kidney) from *Mus caroli* and *Mus pahari* using Qiagen’s RNeasy kit following the manufacturer’s instructions. RNA-seq libraries were generated with Illumina’s TruSeq Ribo-Zero strand specific kit and then sequenced on the Illumina HiSeq2000 platform with 100 bp paired-end reads. The annotation of the *Abp* gene clusters was refined with a combination of BLAST (Altschul et al. 1990), hmmsearch (Finn et al. 2011) and exonerate (Slater and Birney 2005). The relationship between the *Scgb* and *Abp* nomenclatures is described earlier. See **Supplementary Material sections SM1.6 and SM4.5** for more details.

### Divergence time estimation

The divergence times of *Mus musculus* from *Mus caroli* and *Mus pahari* was estimated based on a set of four-fold degenerate sites from amino acids conserved across all mammals. Three different subsets of four-fold degenerate sites with similar size were created based on: (i) random selection; (ii) tissues-specific genes; (iii) housekeeping genes. BEAST 2 (Bouckaert et al. 2014) was used to infer the divergence time independently with the three datasets of four-fold degenerate sites and different evolutionary models (calibrated Yule model, Birth–Death Model, GTR, HKY85, strict clock, uncorrelated relaxed clock). Fossil record information of the mouse-rat divergence (Jacobs and Flynn 2005) was used to calibrate the molecular clock in all our analyses. See **Supplementary Material section SM1.16** for more details.

### Chromosome rearrangement analysis

The synteny breaks involving large genomic regions among *Mus musculus*, *Mus caroli* and *Mus pahari* were identified with the reciprocal cross-species chromosome painting experiments described in **Supplementary Material section SM1.3**. To further define the evolutionarily syntenic breakpoints on the chromosomes of the C57BL/6J strain between *Mus musculus* and *Mus pahari*, a Mouse CGH (244k) microarray was used with the chromosome-specific DNA libraries of *Mus pahari.* The Mouse CGH array was analysed using the CGHweb tool (Lai et al. 2008), with default parameters. For the comparison between *Mus musculus* and rat and between all four Hominidaes, inter-chromosomal synteny breaks involving genomic regions longer than 3 MB were identified and selected using the synteny map in Ensembl v82 (Aken et al. 2017).

To estimate the rate of inter-chromosomal rearrangements in each clade, we created a distance matrix based on the number of synteny breaks. The matrix was used to compute a neighborjoining tree. The branch length from the resulting tree represents an estimation of the number of synteny breaks that occurred in the branch **(Figure S1.4D**).

Repeat enrichment in a +/-40 Mb region around the breakpoints was analysed by counting the occurrence of each repeat element in 200 kb sliding windows and averaging over all breakpoints. For each averaged window, a z-score was calculated based on the 80 Mb region analysed (excluding the +/-2 Mb region around the breakpoint). The size of +/-40 Mb was chosen since it is the longest possible region that does not include a start or end of a chromosome.

See **Supplementary Material section SM2** for more details.

### Evolutionary rate analysis

The nucleotide sequence divergence between *Mus musculus* and the other three murid species as well as between human and other Hominidae was estimated from LASTZ pairwise alignments following the Ensembl methodology (Herrero et al. 2016). For each clade and each genomic class, the value of the nucleotide divergence against the divergence time was plotted for each pair of species involved in the comparison. The rate of nucleotide divergence from each clade was derived from a linear regression. An ANCOVA test was used to evaluate the statistical significance of the difference of rates between each genomic category, with the rate as response variable and the genomic category as a fixed factor.

The rate of unshared genomic segments between *Mus musculus* and other Muridae as well as between human and other Hominidae was estimated from LASTZ pairwise alignments as defined above. A genomic region was defined as shared between two species if the region had an alignment between the two species with less than 50% of gapped sequence. For each clade and each genomic class, the value of the unshared genomic segments was plotted against the divergence time for each pair of species involved in the comparison. The turnover of genomic segments from each clade was derived from a linear regression. An ANCOVA test was used for evaluation of the statistical significance of the difference of turnover between each genomic category, again with turnover rate as response and the genomic category as a fixed factor.

See **Supplementary Material section SM3** for more details.

### Repeat analysis

Repeat elements were identified with RepeatMasker 3.2.8 (A.F.A. Smit, R. Hubley & P. Green RepeatMasker at http://repeatmasker.org) using the rodent repeat libraries for the four Muridae genomes and the primate repeat library for the four Hominidae genomes. Simple repeats and microsatellite elements were removed. Fragmented hits identified by RepeatMasker as belonging to a same repeat were merged. The age of each repeat element was estimated as

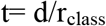

where d is the sequence identity of the repeat with its consensus sequence and rclass is the nucleotide evolutionary rate of the repeat class. The rate was calculated from the ancestral repeats (i.e repeated elements shared between the four Muridae or the four Hominidae genomes). See **Supplementary Material section SM4** for more details.

### Retrocopy analysis

Retrocopies in the Muridae and Hominidae genomes were detected as previously described (Navarro and Galante 2013). In order to comprehensively annotate retrocopies in *Mus musculus* and *Homo sapiens* we used a combination of manual and automatic curation workflows. We considered the manually-annotated processed pseudogenes from GENCODE M13 and v24 respectively Pei, 2012 #132} and processed pseudogenes from pseudopipe (Zhang et al. 2006; Sisu et al. 2014)). Mature transcript sequences were derived from Ensembl v86 and aligned to the corresponding reference genome using BLAT (mask=lower;-tileSize=12;-minIdentity=75;-minScore=100). The age of each retrocopy was estimated as

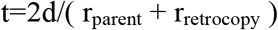

where d is the sequence identity between a retrocopy and its parental gene; rparent is the nucleotide evolutionary rate of the parental gene defined from the set of one-to-one gene-orthologues shared between the four Muridae or four Hominidae; rretrocopy is the nucleotide evolutionary rate of the retrocopies calculated from the retrocopies shared between the four Muridae or the four Hominidae). See **Supplementary Material section SM4** for more details.

### CTCF binding site analysis

We profiled the binding of CTCF in livers of *Mus musculus* C57BL/6J, *Mus caroli* CAROLI/EiJ*, Mus pahari*/EiJ and *Rattus norvegicus* using the ChIP-seq protocol described in Schmidt et al. (Schmidt et al. 2009). The paired-end libraries were sequenced at 100 bp on the HiSeq2000 platform. In addition, the dataset from Schwalie et al (Schwalie et al. 2013) was used to identify the CTCF binding sites in primates. Sequencing reads were aligned to the appropriate reference genome using Bowtie 2 version 2.2.6 (Langmead and Salzberg 2012). MACS version 1.4.2 ((Zhang et al. 2008) was used with a p-value threshold of 0.001 to call read enrichment representing CTCF binding sites. Peaks present in at least two biological replicates were used for the analysis. The binding motif in each CTCF binding region was identified with the FIMO program from the MEME suite version 4.10.2 (Bailey et al. 2015) and using the CTCF position weight matrix (CTCF.p2) from the SwissRegulon database (Pachkov et al. 2013). See **Supplementary Material sections SM1.11 and SM4** for more details.

### SINE B2_Mm1 neighbor-joining classification

SINE B2_Mm1 sequences from the three *Mus* species were selected after filtering out sequences with the following characteristics (i) shorter than 150 bp; (ii) at least one unknown nucleotide (N); and (iii) more than 10% of substitution, insertion or deletion with the SINE B2_Mm1 consensus sequence. The sequences were aligned using MAFFT version 7.222 (Katoh and Standley 2013) and the alignment was used to calculate a neighbor-joining tree using FastTree version 2.1.9 with local bootstrap and minimum-evolution model. The ancestral sequence of the B2_Mm1 CTCF binding motif was inferred using FASTML (Ashkenazy et al. 2012), with the neighbor-joining method and the JC model. A second independent approach based on PRANK (Loytynoja and Goldman 2010), with the options -showanc -keep –njtree was used to confirm the ancestral sequence inference. See **Supplementary Material sections SM5.5-SM7** for more details.

## Data Access

The genome assemblies of *Mus caroli* and *Mus pahari* were submitted to the European Nucleotide Archive (www.ebi.ac.uk/ena) and are available with accession numbers <u>GCA_900094665</u> (*Mus caroli*) and <u>GCA_900095145</u> (*Mus pahari*). All reads from the ChIP-seq and RNA-seq experiments in this study were submitted to ArrayExpress (www.ebi.ac.uk/arrayexpress) and are available with accession numbers E-MTAB-5768 (RNA-seq) and E-MTAB-5769 (ChIP-seq). A supplemental web page with links to raw data and other information is available at http://www.ebi.ac.uk/research/flicek/publications/FOG21.

## Author Contributions

Study design, project leadership and manuscript writing: DT, DTO, PF; genome sequencing and assembly: IS, MK, DT, AD, SA, KS, AZ, MD, WC, LJ, LG, SP, KH, MG, LC, TK; comparative genomics and genome annotation: IF, MS, BP, CC, MM, WA, BA, FM; evolutionary analysis: DM-G, DT, A-AJ, VC; introgression analysis: CVO, BW; chromosome rearrangements analysis: DT, FY; repeat analysis: DT; retrocopy analysis: FCPN, DT, CS, MG; CTCF and repeat analysis: MR, CF, DT, MH; Abp region analysis: VJ, GY, RCK, CML; reagent supply: FV, DA, AB,

## Acknowledgements

This project was supported by the Wellcome Trust (grant numbers WT108749/Z/15/Z, WT098051, WT202878/Z/16/Z and WT202878/B/16/Z) the National Human Genome Research Institute (U41HG007234), Cancer Research UK (20412), the European Research Council (615584), and the European Molecular Biology Laboratory. The research leading to these results has received funding from the European Community’s Seventh Framework Programme (FP7/2010-2014) under grant agreement 244356 (NextGen). The research leading to these results has received funding from the European Union’s Seventh Framework Programme (FP7/2007-2013) under grant agreement HEALTH-F4-2010-241504 (EURATRANS).

We thank the genomics, bioinformatics, and BRU cores at the CRUK Cambridge Institute for technical support, the sequencing facilities at the Wellcome Trust Sanger Institute and computational support from EMBL-EBI and WTSI as well as the Conservatoire Génétique de la Souris Sauvage (ISEM, France) and Plateforme Cytogénomique évolutive of the LabEx CeMEB. We also thanks Bee Ling N, Beiyuan Fu, and Sandra Louzada, Mark Simmonds for assistance in chromosome sorting, chromosome painting, and array painting.

